# Cortical thickness in right frontal and left lingual gyri differentially mediate episodic memory for spatial contextual details across the adult lifespan

**DOI:** 10.1101/289447

**Authors:** A. Swierkot, M. N. Rajah

## Abstract

Healthy aging is associated with declines in episodic memory and with widespread cortical thinning. These parallel declines suggest that age-related changes in cortical thickness may contribute to episodic memory decline with age. The current study uses a cross-sectional study design to examine whether regional cortical thickness mediates the relationship between age and episodic memory, as measured by a context memory task for faces. Mediation and conditional mediation models were tested using bootstrapping in order to determine how age-associated changes in regional cortical thickness mediated age-associated changes in performance on the context memory task. We observed that right superior frontal cortical thickness conditionally mediated spatial context memory only in middle-aged and older adults; and right caudal middle frontal cortical thickness conditionally mediated context memory only in older adults. Left lingual cortical thickness mediated spatial context memory across the adult lifespan, but this effect was most evident at midlife. Right parahippocampal cortical thickness mediated context memory, independent of age. We conclude that our cortical thickness results were generally consistent with the posterior-to-anterior shift in aging hypothesis (Davis et al., 2008) for episodic memory.

## 1. Introduction

Healthy aging is associated with declines in the ability to remember an item or event in rich contextual detail (episodic memory; Tulving, 1984). Studies have shown that episodic memory tasks that assess one’s ability to encode and retrieve contextual details (item-context associative memory tasks) are more sensitive at detecting age-associated episodic memory deficits, compared to item recognition tasks (Cansino, 2009; Dulas & Duarte, 2012; Glisky, Polster, & Routhieaux, 1995; Mattson, Wang, de Chastelaine, & Rugg, 2013; Rajah, Languay, & Valiquette, 2010; Schacter, Kaszniak, Kihlstrom, & Valdiserri, 1991; Trott, Friedman, Ritter, & Fabiani, 1997). FMRI studies of young adults have shown increased activity in medial temporal lobes (MTL) and prefrontal cortices (PFC) during associative context memory tasks, compared to item recognition tasks (Davachi, 2006; Maillet & Rajah, 2013; Shing et al., 2010; Staresina & Davachi, 2008). Moreover, it has been hypothesized that successful encoding and retrieval of item-context associations in young adults relies on the engagement of a distributed network of brain regions including sensory cortices, MTL, inferior parietal cortex (IPC) and PFC (H. Kim, Daselaar, & Cabeza, 2010; Nyberg et al., 2000; Rugg & Vilberg, 2013). Thus, it follows that age-related decrements in item-context associative memory tasks may be mediated by structural changes in any (or all) of the brain regions implicated with successful memory encoding and retrieval in young adults (Braskie, Small, & Bookheimer, 2009; Spencer & Raz, 1994).

Several studies have reported age-related volumetric decline in brain regions related to encoding and/or retrieval of contextual details from episodic memory (Fjell et al., 2013; Hogstrom, Westlye, Walhovd, & Fjell, 2013; Raz, Ghisletta, Rodrigue, Kennedy, & Lindenberger, 2010; Walhovd et al., 2011). Salat et al (2004) reported reductions in cortical thickness with age in PFC, supramarginal gyrus, and calcarine sulcus, with relative preservation of lateral temporal cortex thickness. Raz et al (2005) also reported a steady decline in PFC grey matter volumes with age, starting in young adulthood. However, five year longitudinal follow-up data indicated there was notable individual variability in how PFC volumes changed with age, with some subjects exhibiting volumetric decline, and other subjects exhibiting either no change or increased PFC volume at five year follow-up (Raz et al., 2005). In contrast to a general decrease in PFC volumes throughout adulthood, hippocampal volume decline with age exhibits a non-linear pattern. For example, Walhovd et al (2011) have reported changes in hippocampal volumes with age resembling an inverted-U shape, which was suggestive of accelerated volume loss in this region in late midlife onwards. This observation was corroborated by Fjell et al (2013). Therefore, there is significant evidence that aging is related to structural decline in several brain regions associated with episodic memory.

However, fewer studies have examined whether age-related declines in brain volume and/or cortical thickness directly relates to age-related decline in episodic memory tasks. In other words, do age-related differences in regional cortical thickness or brain volume mediate the effect of age on memory function? In one study, Walhovd et al. (2006) used a cross-sectional study design to examine the relationship between cortical thickness and episodic memory, as measured by long-term recall on the California Verbal Learning Task (CVLT), in a full lifespan sample. Through a whole-brain vertex-wise analysis they found that *after controlling for age*, long-term recall was positively associated with cortical thickness in bilateral gyrus rectus, bilateral medial frontal gyrus, bilateral parieto-occipital sulcus, bilateral lingual gyri, right temporal and parietal lobes, and left precuneus throughout the adult lifespan (Walhovd et al., 2006). Importantly, this analysis uncovered these brain-behaviour relationships after controlling for age, meaning that this analysis explained how individual differences in cortical thickness related to memory. However, it did not assess whether *cortical thickness* mediated the effect of age on memory performance. Head, Rodrigue, Kennedy, & Raz (2008) aimed to answer this question by conducting a path analysis to test the hypothesis that age-associated changes in cognitive function was mediated by changes in brain volumes. They found that mean PFC volume mediated age-associated variance in episodic memory performance via its role in executive processing tasks (Head, Rodrigue, Kennedy, & Raz, 2008). However, this analysis did not consider the possibility that brain-behaviour relationships may change with age. For instance, there may be a relationship between cortical thickness or volume in one brain region and behaviour during one period of the lifespan, and not another; indicative of non-linear changes in brain-behavior associations with age. In other words, it is possible that due to the different linear and non-linear patterns of structural decline in distinct brain regions with age (Walhovd et al., 2011), that not only could structural decline mediate age-related differences in episodic memory, but that this brain-behavior relationship could be different at different ages (moderated by age). For example, it could be that in younger adulthood item-context associative memory performance may be more directly predictive of cortical thickness or volume in MTL, but with older age performance may more related to cortical thickness in PFC. For example, we previously found that anterior HC volume was positively associated to context memory performance in young adults (YA), but not older adults (OA) (Rajah, Kromas, Han, & Pruessner, 2010). In that study we hypothesized that due to the hippocampal decline observed in OA compared to YA, that memory performance in OA may have been dependent on more anterior cortical regions, such as the PFC, as predicted by the posterior-to-anterior shifts in aging (PASA) model of cognitive aging (Davis, Dennis, Daselaar, Fleck, & Cabeza, 2007). However, to our knowledge no study has examined how age-related declines in item-context associative memory may be mediated by regional changes in cortical thickness, and if this mediation could be moderated by age. We aimed to directly test this hypothesis in our current study.

In the current study high-resolution structural MRIs were obtained from 113 healthy young, middle-aged, and older adults who also performed a spatial context memory task. We examined age related differences in cortical thickness across the whole brain, and identified regions of interests (ROIs) that we have previously identified as being implicated in this spatial context memory task (Ankudowich et al., 2016), that also exhibited an age-related difference in cortical thickness. We then built conditional mediation models that simultaneously considered associations between age, retrieval accuracy and cortical thickness; whether cortical thickness mediated the effect of age on retrieval accuracy; and, whether this mediational effect was also moderated by age (conditional mediation). More specifically using the conditional mediation model we simultaneously assess the following five patterns of associations: 1. Age is related to retrieval accuracy from spatial context memory tasks; 2. Age is related to cortical thickness; 3. Cortical thickness is related to retrieval accuracy (after controlling for age), 4. Cortical thickness mediates the relationship between age and spatial context retrieval accuracy; and 5. Age can moderate the relationship between cortical thickness and retrieval accuracy, at specific times/ages (i.e. young, middle-age and older adulthood). Based on previous findings we predicted spatial context retrieval accuracy would decline with age (Ankudowich et al., 2016; Cansino, 2009; Dulas & Duarte, 2011; Glisky, Polster, & Routhieaux, 1995; Mattson, Wang, de Chastelaine, & Rugg, 2013; Rajah, Languay, & Valiquette, 2010; Schacter, Kaszniak, Kihlstrom, & Valdiserri, 1991; Trott, Friedman, Ritter, & Fabiani, 1997). We also predicted that age-related declines in cortical thickness would be present throughout the cortex as observed in prior studies (Allen, Bruss, Brown, & Damasio, 2005; Burzynska et al., 2013; Fjell et al., 2009; Raz et al., 2005; Salat et al., 2004; Thambisetty et al., 2010). Finally, based on our prior findings (Rajah, Kromas, et al., 2010) and those of others (Davis, Dennis, Daselaar, Fleck, & Cabeza, 2008), we hypothesized that cortical thickness in posterior cortical regions would be positively associated with spatial context retrieval accuracy in young adults, and less so with advanced age. We also predicted that cortical thickness in prefrontal regions would be positively associated spatial context retrieval accuracy in older adults, and less so in younger adulthood. Therefore, we predicted that consistent with the PASA model of cognitive aging (Davis et al., 2008), age would moderate the effect of cortical thickness on spatial context memory in posterior cortex and PFC.

## 2. Methods

### Participants

One hundred and thirteen adults (age range 19-76 years; mean age 46.92; mean education 15.69 years; 77 females) participated in the study. At the time of testing, all participants were healthy and had no history of neurological or psychological illness. All participants were right-handed as measured by the Edinburgh Inventory for Handedness (Oldfield, 1971). To screen out individuals suffering from psychiatric symptoms and dementia, and to obtain measures of memory and language function, volunteers were administered the following battery of neuropsychological tests (Kwon et al., 2016): the Mini-International Neuropsychiatric Interview (M.I.N.I.) [inclusion cut-off score ≤ 2, (Sheehan et al., 1998)], Montreal Cognitive Assessment [MOCA, exclusion cut-off score < 27, (Nasreddine et al., 2005)], the Beck Depression Inventory (BDI) [inclusion cut-off < 15 (Beck & Steer, 1987)], the California Verbal Learning Task (CVLT) [exclusion cut off determined per case using age & education (Norman, Evans, Miller, & Heaton, 2000)], the American National Adult Reading Test (NART) [inclusion cut-off ≤ 2.5 SD (Spreen & Strauss, 1998)]. Additional medical exclusion criteria included having a history of or current diagnosis of diabetes, untreated cataracts and glaucoma, and a current diagnosis of high cholesterol levels and/or high blood pressure left untreated in the past 2 years. Moreover, anyone with a first-degree relative who had been diagnosed with Alzheimer’s disease was excluded from the study. Table 1 provides the demographic characteristics of the sample tested in the current study.

**Table 1:** Demographics and Behavioral Task Performance

| GROUP |  | AGE | EDU | CVLT-FR | CVLT-CR | CVLT-RG | MOCA | SE<br>accuracy | SH<br>accuracy | SE<br>RT(ms) | SH RT<br>(ms) |
| --- | --- | --- | --- | --- | --- | --- | --- | --- | --- | --- | --- |
| **Young**<br>**Adults** | Mean | 26.18 | 16.10 | 13.85 | 13.98 | 15.50 | 29.30 | 0.88 | 0.87 | 2241.87 | 2351.86 |
|  | N | 40 | 40 | 40 | 40 | 40 | 40 | 40 | 40 | 40 | 40 |
|  | Std. Dev. | 3.80 | 1.88 | 1.75 | 1.73 | 0.72 | 0.97 | 0.09 | 0.10 | 527.10 | 498.13 |
| **Middle-**<br>**Aged**<br>**Adults** | Mean | 48.88 | 15.39 | 12.67 | 13.27 | 15.33 | 28.58 | 0.84 | 0.78 | 2554.95 | 2675.70 |
|  | N | 33 | 33 | 33 | 33 | 33 | 33 | 33 | 33 | 33 | 33 |
|  | Std. Dev. | 5.77 | 2.12 | 2.31 | 1.99 | 0.74 | 1.28 | 0.10 | 0.13 | 499.51 | 421.45 |
| **Older**<br>**Adults** | Mean | 66.55 | 15.53 | 12.88 | 13.15 | 15.15 | 28.73 | 0.82 | 0.77 | 2784.87 | 2859.99 |
|  | N | 40 | 40 | 40 | 40 | 40 | 40 | 40 | 40 | 40 | 40 |
|  | Std. Dev. | 3.60 | 2.20 | 2.43 | 2.08 | 0.89 | 1.20 | 0.10 | 0.11 | 470.46 | 475.10 |
| **Total** | Mean | 47.10 | 15.69 | 13.16 | 13.48 | 15.33 | 28.88 | 0.85 | 0.81 | 2525.51 | 2626.30 |
|  | N | 113 | 113 | 113 | 113 | 113 | 113 | 113 | 113 | 113 | 113 |
|  | Std. Dev. | 17.65 | 2.07 | 2.22 | 1.96 | 0.80 | 1.18 | 0.10 | 0.12 | 546.00 | 512.71 |

**Note:** The mean and standard deviation (Std. Dev) for the sample (N) analysed. EDU = education (yrs), CVLT = California Verbal Learning Task, delayed accuracy, FR = delayed free recall, CR = delayed cued recall, RG = delayed recognition, MOCA = Montreal Cognitive Assessment, SE = spatial easy task, SH = spatial hard task, RT = reaction time.

### Behavioural methods

The current study is part of a larger fMRI study exploring the structural and functional neural correlates of context memory across the adult lifespan (E. Ankudowich, Pasvanis, & Rajah, 2017; E. Ankudowich, Pasvanis, S., Rajah, M. N., 2016). In the current study we focus on the association between age, cortical volume and spatial context memory for faces. In this paradigm, subjects were told that they would be participating in a computer-based memory experiment for non-famous faces and would be required to memorize and later remember the left/right spatial location of faces. Subjects then performed either easy or hard versions of a left/right spatial context memory tasks. The rationale for the easy/hard manipulation was to allow for the examination of age effect, performance effects and age* performance interactions in the published fMRI data (Kwon et al., 2016; Ankudowich et al, 2016).

At encoding, subjects were presented with either 6 black and white photographs of human faces (easy version) or 12 faces (hard version) to memorize. Faces were presented one at a time (2 secs per face), on either the left side or the right side of a central fixation cross, on a computer screen. Subjects were required to memorize the face and it’s left/right spatial location, and also rate each face as either pleasant/neutral. The pleasantness rating was used improve memory success in the current paradigm since prior research has shown that making such social-emotional evaluation of stimuli at encoding was associated with improved memory at retrieval (Grady, Bernstein, Beig, & Siegenthaler, 2002). There was a variable inter-trial interval (ITI; 2.2, 4.4, or 8.8 sec).

Following encoding there was a 1-minute distraction task during which subjects were presented with 5 sets of 2 words and asked to put them in alphabetical order. The distraction task was used to prevent rehearsal of face stimuli and ensure retrieval was from episodic memory, not working memory. At retrieval, participants were shown vertically oriented pairs of previously viewed faces. Each retrieval pair was presented for 6 seconds with variable ITI. Upon viewing a pair, participants were instructed to select which face they previously saw on either the left or the right-side of the computer screen, depending on the instruction cue. Therefore, a “correct retrieval” response required the recollection of the face-location association of either both stimuli, or just one of the two stimuli. As such, subjects could have employed a recall-to-reject strategy to accurately perform this task. Alternatively, subjects could have guessed during retrieval (chance = 50%). However, all subjects in this study performed above chance. Subjects performed 12 spatial easy (SE) memory tasks and 6 spatial hard (SH) memory tasks in total. There were 72 encoding trials and 36 retrieval trials per task type.

### MRI data acquisition

Scanning of subjects was performed on a 3T Siemens Trio scanner at the Douglas Brain Imaging Center. Subjects were asked to lie in a supine position in the MRI scanner while wearing a standard head coil. At the start of the experiment, a high-resolution T1-weighted anatomical images was acquired from each subject using a 3D gradient echo MPRAGE sequence (TR = 2300 msec, TE = 2.98 msec, flip angle 9°, 176 1mm sagittal slices, 1×1×1 mm voxels, FOV 256mm^2^).

### Statistical analyses

#### Behavioral Analysis

To address concerns of education as a confounding variable, age was regressed against education to determine whether they were significantly related. Accuracy on the spatial tasks was calculated as a percentage of correct responses on easy and hard tasks separately. Two simple linear regressions were run to examine age-related changes in spatial context memory. The two behavioural metrics (spatial easy accuracy and spatial hard accuracy) were regressed against age to examine age-associated variability in context memory within the current sample.

#### MRI Analyses

##### Cortical Thickness Analysis

Cortical thickness was estimated at 81924 points across the cortex (40962 per hemisphere) on the T1-weighted images using the automated CIVET 1.1.10 pipeline (Collins, Neelin, Peters, & Evans, 1994; Fahim, Yoon, Sandor, Frey, & Evans, 2009; Rajaprakash, Chakravarty, Lerch, & Rovet, 2014). Each subjects’ T1 image was linearly registered to the standardized MNI ICBM 152 model using a 9-parameter linear transformation (Mazziotta et al., 2001) and corrected for intensity non-uniformity using the N3 algorithm (Sled, Zijdenbos, & Evans, 1998). A non-linear registration was then applied. Next, images were segmented into white matter (WM), grey matter, cerebral spinal fluid (CSF) and background (Zijdenbos, Forghani, & Evans, 2002). Partial volume estimates were applied to define deep sulci (J. S. Kim et al., 2005). Deformable ellipsoid polygonal models were optimized using Constrained Laplacian Automated Segmentation with Proximities (CLASP) to fit the WM-grey matter and grey matter-CSF boundaries (J. S. Kim et al., 2005; MacDonald, Kabani, Avis, & Evans, 2000). Cortical thickness was calculated by measuring the distance between the white and grey surfaces (Lerch, 2001; Lerch & Evans, 2005). These measurements were then blurred using a 20mm kernel to produce the final thickness maps.

To examine relationships between cortical thickness and age in the current dataset, a whole brain vertex-wise analysis of cortical thickness changes with age was conducted, examining left and right hemispheres separately, as described above. The following general linear model (GLM) was fitted to each one of the 40962 cortical points per hemisphere:

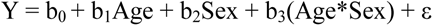

False discovery rate (FDR, p<0.05) corrections were applied to p-values to account for multiple comparisons.

##### Selection of ROIs

Next, to examine relationships between age-related cortical thickness and spatial context memory, we selected regions of interest (ROIs) from the LONI Probabilistic Brain Atlas (Shattuck et al., 2009) that showed cortical thinning with age, based on our cortical thickness analysis. We used an automated tool (https://github.com/CobraLab/lpba40-local-civet-measures) to obtain mean cortical thickness for these ROIs for each hemisphere.

##### Mediation and Conditional Mediation Models

The goal of the current study was to determine if the effect of age on spatial context retrieval accuracy was *mediated* by the cortical thickness of specific ROIs (that also exhibited age-related differences in thickness), ***and*** if this mediational effect was contingent on specific stages of adult development (i.e. YA, MA or OA; *moderated or conditional mediation* by age-group). In other words, we wanted to examine both linear mediational effects – whether ROI cortical thickness mediates age-related changes in retrieval accuracy; and non-linear mediational effect - whether ROI cortical thickness mediates age-related changes in retrieval accuracy, at specific stages of adult development (i.e. young adulthood, middle-age and older age). To examine conditional mediation effects, we needed to use a method that considered the following 5 key relationships present in our lifespan dataset, across multiple ROIs *simultaneously*: 1. Age is directly related to spatial context retrieval accuracy; 2. Age is directly related to cortical thickness; 3. Cortical thickness is directly related to retrieval accuracy (after controlling for age); 4. Cortical thickness mediates the relationship between age and retrieval accuracy; and 5. Age moderates the relationship between cortical thickness and spatial context memory performance. The parallel mediation (Model 4) model from the SPSS custom dialogue tool PROCESS (version 2.16) (Hayes & Rockwood, 2017; Preacher & Hayes, 2004, 2008) addresses the first four relationships. The conditional mediation model (Model 74) in PROCESS addresses the first three and fifth relationships while simultaneously examining how cortical thickness mediates age-related differences in memory performance at specific stages in adulthood (young, middle-aged and old)— conditional (or moderated) mediation. The statistical diagram for these models are presented in Figure 1. In line with the vertex-wise analysis, separate models were applied for the left and right hemispheres.

**Figure 1.**
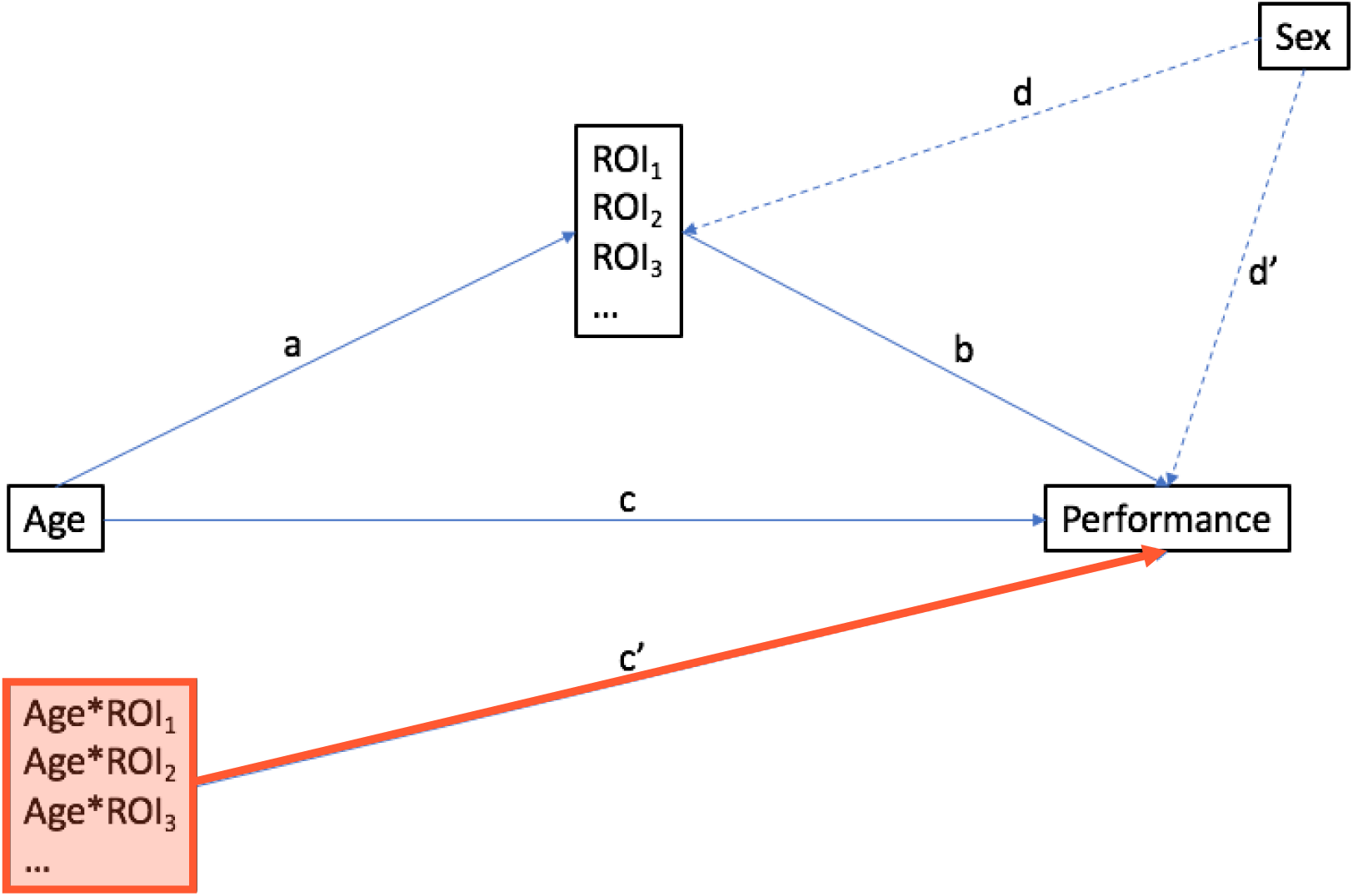
Statistical Diagram of Mediation and Conditional Mediation Models tested. The unfilled boxes and blue lines reflect the mediation model (model 4) from PROCESS (Hayes, 2013). The direct effect of age on memory performance is reflected by the path c and the indirect (mediational) effect of cortical thickness of all 9 ROIs for left or right hemisphere models respectively are considered simultaneously in parallel, and calculated as follows: M_i_ = a_i_b_i_. The conditional mediation model (model 74) is model 4 plus the Age*ROI moderated mediational effect tested at specific quintiles, coloured in red. Conditional indirect effect of age on performance via cortical thickness is calculated as M_i_=a_i_(b_i_+ c_i_’X) and the direct effect of age on performance is calculated as X = c + c_i_’M. Sex is a covariate of no interest in the models. Path a reflects age→ROI coefficient; b reflects ROI→Performance coefficient; c reflects age→performance coefficient; c’ reflects Age*ROI→Performance coefficient; d reflects Sex→ROI coefficient; d’ reflects Sex→Performance coefficient.

Model 4, as diagrammed in Figure 1, simultaneously tests for mediation effects in all ROIs. In this model age was the independent variable and SE or SH retrieval accuracy was the dependent variable, and the mediators were the left and right hemisphere ROIs identified from the cortical thickness analysis, respectively. Model 74, as diagrammed in Figure 1, is exactly the same as Model 4, except that it also considered age as a moderator (age*ROI) so that it tested for conditional mediation at specific points in the lifespan. The specific points were selected based on the 10^th^, 25^th^, 50^th^, 75^th^ and 90^th^ percentiles of the moderator (age) distribution. In the current study these quantiles matched the following ages: 24 yrs, 27 yrs, 48 yrs, 64 yrs and 68 yrs, respectively. In conducting these mediational analyses PROCESS also simultaneously conducts traditional regression analyses. It should be noted that the regression coefficients and mediation effects are related, but distinct. In other words, significant regression coefficients do not necessarily mean that significant moderated mediation is present and vice versa. Separate analyses were run for left and right hemispheres and for spatial easy and spatial hard tasks, for a total of 4 models: 1) SE - Right hemisphere ROI model; 2) SE - Left hemisphere ROI model; 3) SH - Right hemisphere ROI model; and 4) SH - Left hemisphere ROI model. Significance was assessed at p<0.05. PROCESS mean-centers the variables in the analysis to mitigate concerns of multi-collinearity.

#### Testing significant direct effects with regression

The first key relationships of interest reflect direct regression effects, the significance of which can be determined by examining the associated regression coefficient outputs from PROCESS (also known as weights). The age→performance coefficient represented *age-associated* variance in performance— how performance varied with age. The age→ROI coefficient represented *age-associated* variance in regional cortical thickness— how cortical thickness varied with age. The ROI→performance coefficient represented *cortical thickness-associated* variance in performance *homogeneously* across the entire lifespan sample—how cortical thickness was linearly related to performance, *after controlling for age*. In other words, a significant ROI→performance coefficient represented how a region was related to performance after variability due to age was removed from the ROI and performance variables. This kind of relationship is age-independent.

#### Testing significant moderation with regression

To test whether age moderated the relationship between cortical thickness and performance, age*ROI terms were included in our conditional mediation models (model 74; see figure 1). The age*ROI→performance coefficient represented *cortical thickness-associated* variance in performance *differentially* across the lifespan sample—how cortical thickness was non-linearly related to performance.

#### Testing the significance of mediational and conditional mediation effects

The significance of mediation of age-related differences in retrieval accuracy by cortical thickness in specific ROIs; and moderated mediation at different stages in adulthood was assessed using bootstrapping. All ROIs included in the model were simultaneously evaluated, negating the need for multiple comparisons. Significance was evaluated by calculating bias-corrected 95% confidence intervals using bootstrapping with 1000 samples. Sex was included as a covariate of no interest in these analyses. Conditional mediation was probed at the quantiles described above. A significant mediation and/or conditional mediation effect was represented as a bootstrap confidence interval entirely above, or entirely below zero. A significant mediation effect indicated that age-related differences in retrieval accuracy were mediated by cortical thickness in specific ROIs, across the adult lifespan (a linear effect). A significant conditional mediation effect indicated that cortical thickness in specific ROI mediated retrieval accuracy at specific points in the adult lifespan, as defined by the quantiles. This analysis helped identify non-linear mediational each of ROI cortical thickness on retrieval accuracy in the adult lifespan.

## 3. Results

### 3.1. Age and education

Table 1 presents the demographics for the sample. Mitigating concerns of education confounding age-effects in our analyses, years of education was regressed against age, and it was found that age did not significantly predict education in our sample (F(1,112)=3.583, p=0.061; adjusted *R^2^*=0.022).

### 3.2. Age-related changes in spatial context accuracy

Our simple linear regression analysis uncovered that both behavioural metrics were significantly related to age. Spatial easy accuracy decreased with age (F(1,112)= 9.01, p = 0.003; Adjusted *R^2^*=0.066), and spatial hard accuracy decreased with age (F(1,112) = 18.92, p < 0.001; Adjusted *R^2^*=0.137). Mean values for SE and SH accuracy and retrieval are presented in Table 1.

### 3.3. Age-related cortical thickness changes

Figure 2 shows the results from the vertex-wise cortical thickness analysis testing for the association between cortical thickness and age, while controlling for sex and age-by-sex interactions. This figure illustrates there was a significant distributed pattern of age-associated difference in cortical thickness in frontal, parietal, temporal, and visual regions. Based on the cortical thickness results, nine LONI ROIs were identified in the right hemisphere were: parahippocampal, rectus, superior temporal, middle temporal, angular, supramarginal, superior frontal, caudal middle frontal, and inferior frontal gyri. Nine LONI ROIs were also identified in the left hemisphere were: parahippocampal, rectus, lingual, fusiform, angular, supramarginal, superior frontal, caudal middle frontal, and inferior frontal gyri. These ROIs were the mediators employed in the right and left hemisphere mediation and conditional mediation models tested.

**Figure 2.**
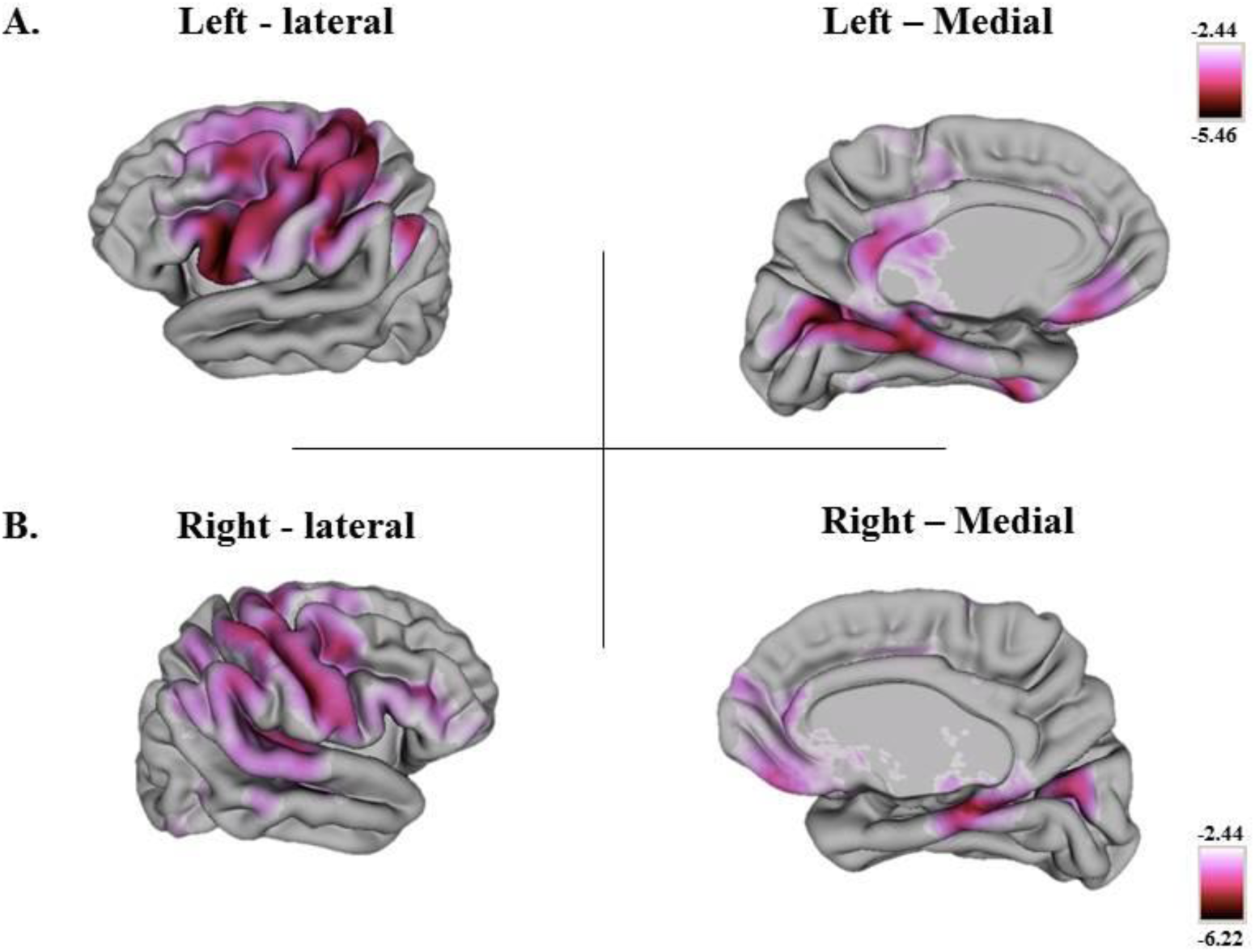
The effect of age on cortical thickness analysis for right (A) and left (B) hemispheres, as measured by CIVET. A general linear model was fitted to 40962 cortical points per hemisphere. This model examined the effect of age on cortical thickness, while taking into account the effects of sex and age-by-sex interactions (see methods for equation). FDR corrections (p<0.05) were applied to the p-values to account for multiple comparisons. The current figure depicts t-maps for the results of this analysis. A threshold was set on the t-maps such that points whose t-values with corresponding corrected p-values less than 0.05 appear in magenta. It should be noted that all significant t-values were negative on both hemispheres, indicating a negative relationship between age and cortical thickness at all points highlighted in magenta

### 3.4. Mediation & Conditional Mediation analyses

Our results indicated that only the mediation models examining the associations between age, ROI cortical thickness and retrieval accuracy during hard spatial context (SH) memory tasks were significant.

#### Right hemisphere models for SH retrieval accuracy

Consistent with the CIVET results, there were significant age-associated differences in cortical thickness in all 9 selected ROIs (age → ROI) observed in both mediational and conditional mediational models. Older adults exhibited significantly thinner cortex compared to younger adults.

##### Model 4: Mediation Model

The overall mediation model for SH retrieval was significant (F(11,101) = 2.78, p =0.003), and was due to SH retrieval accuracy being significantly related to Age (t = −2.60, p = 0.01). However, cortical thickness in the right hemisphere ROIs did not significantly mediate this effect (p > 0.05).

##### Model 74: Conditional Mediation Model

The overall conditional mediation model for SH retrieval accuracy was significant (F(20,92) = 2.27 p = 0.005; Supplementary Table 1S contains the path coefficients for the model). There was a significant conditional mediation of right SFG cortical thickness on SH retrieval accuracy in middle aged and older adults, and a significant conditional mediation of right caudal middle frontal gyrus (MFG) on SH retrieval accuracy in older adults. The bootstrap results for the conditional mediation effect indicated that right SFG cortical thickness did not mediate memory performance at age 24 and 27 yrs, but did significantly mediate SH retrieval accuracy at 48 (bootstrap results, Lower CI to Upper CI: −0.004 to −0.0002), 64 (bootstrap results, Lower CI to Upper CI: −0.007 to −0.001) and 68 (bootstrap results, Lower CI to Upper CI: −0.008 to −0.001) years of age. Right MFG cortical thickness significantly mediated SH retrieval accuracy at age 64 (bootstrap results, Lower CI to Upper CI: to 0.0003 to 0.005) and 68 (bootstrap results, Lower CI to Upper CI: 0.0004 to 0.006). Figure 3 presents scatterplots showing the association between age and right SFG cortical thickness (3A, i), the association between cortical thickness and SH retrieval accuracy across all ages (3B, i), the association between cortical thickness and SH retrieval accuracy in middle aged adults (aged 40 – 59 yrs) and older adults (aged 60 – 78 yrs; 3C, i). Similar scatterplots are presented for right MFG in Figure 3A (ii), 3B(ii) and 3C(ii). Taken together, the results indicate that right frontal cortical thickness mediates the effect of age on SH retrieval accuracy at later life. More specifically, in middle and older aged adults there was a positive mediation between right SFG cortical thickness and SH retrieval accuracy; but, in older age there was also a negative mediation between right MFG cortical thickness and SH retrieval accuracy. In other words, having a thicker right SFG cortex, and a thinner right MFG cortex in older age predicts better SH retrieval accuracy. To directly test this we conducted a post-hoc linear regression analysis. We standardized the dependent variable (SH retrieval accuracy) and all predictor variables: age, right SFG cortical thickness, and right MFG cortical thickness. The results indicated the overall model was trending towards significant (F (3, 36) = 2.27, p =0.097). Yet, importantly, negative beta coefficients were reported for age (standardized beta = −0.13), and right MFG (standardized beta = − 0.26), but a positive beta coefficient was reported for right SFG (standardized beta = 0.57). This verifies the conditional mediation results and confirms that in older adults, thicker right SFG and thinner right MFG predicted better SH retrieval accuracy.

**Figure 3.**
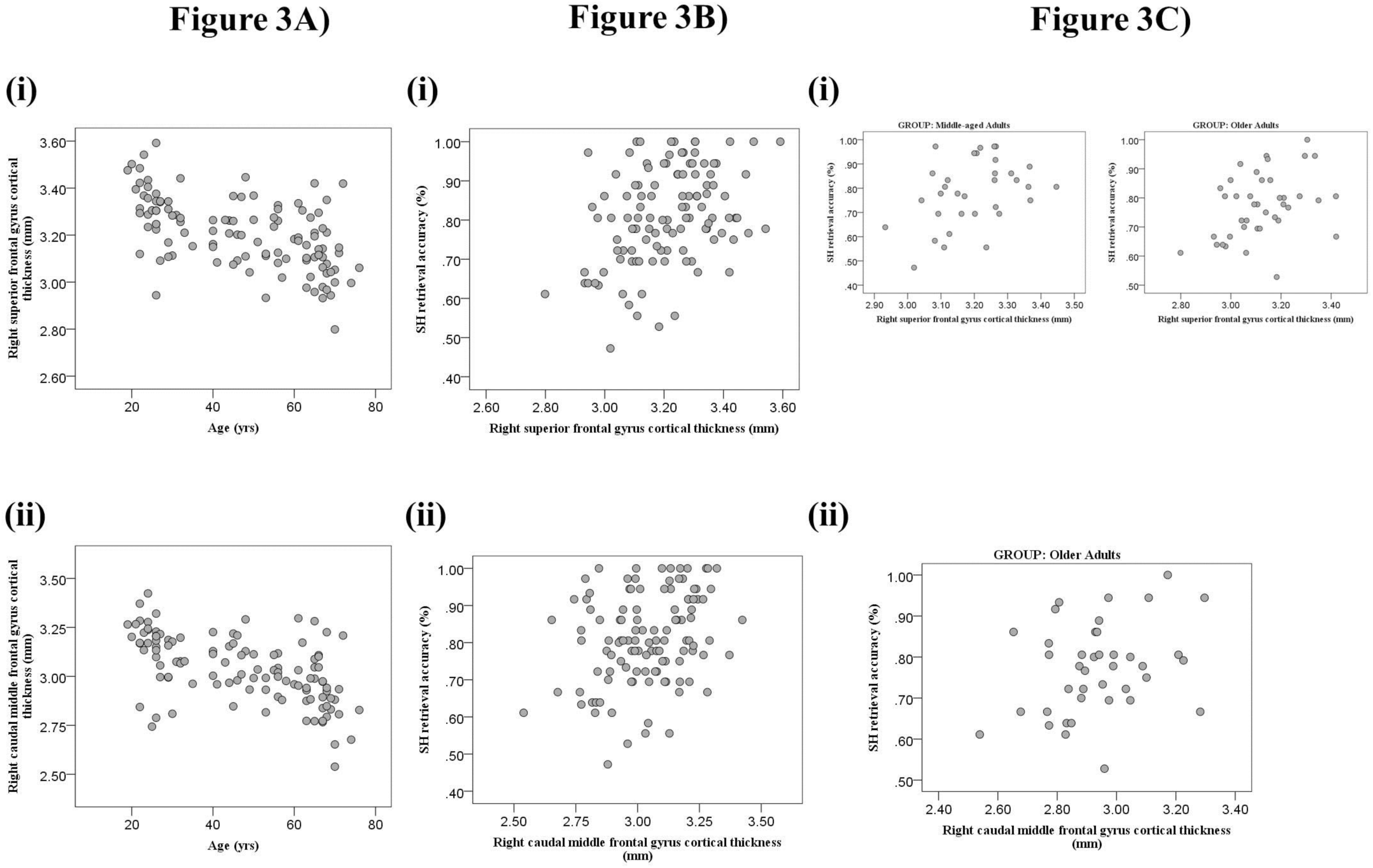
Scatterplots for right frontal ROIs exhibiting significant conditional mediation effects. 3A) association between age and cortical thickness for right SFG (i) and MFG (ii); 3B) the association between cortical thickness and SH retrieval accuracy across all ages for right SFG (i) and MFG (ii); 3C) the association cortical thickness and SH retrieval accuracy for specific age group(s) for which there was a significant conditional mediation effect.

In addition, to the significant conditional mediation effects there was also a near significant direct effect of right PHG cortical thickness → spatial context memory performance (t=1.97, p = 0.052), after controlling for age. Thus, right PHG cortical thickness predicted SH retrieval accuracy, but was not a significant mediator of the effect of age on SH retrieval accuracy. A similar direct effect was observed for right SFG (t = −2.02, p = 0.046), but the significant conditional mediation effect suggests that cortical thickness in this region also mediated the effect of age on SH retrieval accuracy later in life. It should also be noted that after accounting for the conditional mediation effects in the model, the effect of age on SH retrieval accuracy was no longer significant (age, t < 1.0, p > 0.05).

#### Left hemisphere models for SH retrieval

##### Model 4: Mediation Model

The overall mediation model for SH retrieval accuracy was significant (F(11,101) = 3.26, p =0.0008). SH retrieval accuracy was significantly predicted by age (t = −2.14, p = 0.03), and this effect was also significantly mediated by the cortical thickness of left lingual cortex across the adult lifespan (bootstrap results, Lower CI to Upper CI: −0.003 to −0.0004).

##### Model 74: Conditional Mediation Model

The overall conditional mediation model for SH retrieval accuracy was significant (F(20,92) = 2.52 p = 0.002; Supplementary Table 2S contains the path coefficients for the model). Conditional mediation was observed for left lingual gyrus and left angular gyrus. However, no significant direct effects of ROI cortical thickness → performance nor indirect effects of Age*ROI cortical thickness → performance were observed for either ROIs. This indicates that cortical thickness in these ROIs did not predict SH retrieval accuracy nor did age moderate the effect of cortical thickness of these ROIs on SH retrieval accuracy. However, the bootstrap results showed that left lingual gyrus cortical thickness conditionally mediated the effect of age on SH retrieval accuracy at ages 24 (Lower CI to Upper CI: −0.0038 to − 0.0004), 27 (Lower CI to Upper CI: −0.0035 to −0.0005) and 48 (Lower CI to Upper CI: −0.0028 to − 0.0003). Figure 4 presents scatterplots showing the association between age and left lingual gyrus cortical thickness (4A, i), the association between left lingual gyrus cortical thickness and SH retrieval accuracy across all ages (consistent with the aforementioned significant mediation result (4B, i)), and the association between left lingual gyrus cortical thickness and SH retrieval accuracy in young adults (aged 18 –35yrs), and middle aged adults (aged 40 – 59 yrs) (4C, iii). Taken together these results suggest that cortical thickness of left lingual gyrus positively mediated the effect of age on SH retrieval accuracy across the adult lifespan, but this effect is most pronounced at midlife. Interestingly, after combining young adults, we do not see a strong association between left lingual cortical thickness and SH retrieval accuracy in this age group (scatterplot 3C; post-hoc within young adult group ROI→performance regression: F < 1). This implies the significant conditional mediation effect at ages 24 and 27 may reflect an artifact in how the young adult group was split by quintiles.

**Figure 4.**
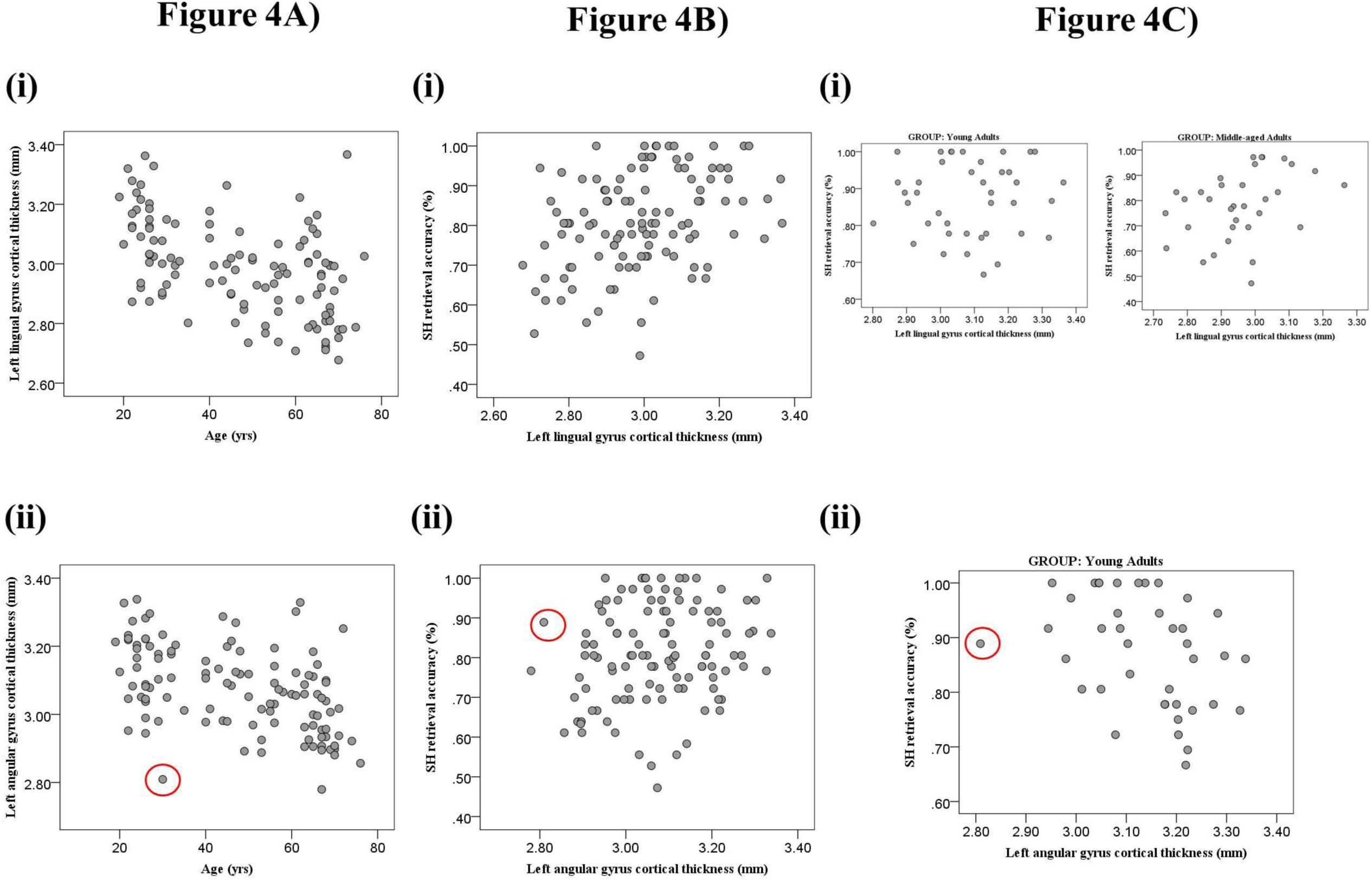
Scatterplots for left hemisphere ROIs exhibiting significant conditional mediation effects. 4A) association between age and cortical thickness for left lingual (i) and angular (ii) gyri; 4B) the association between cortical thickness and SH retrieval accuracy across all ages for left ligual (i) and angular (ii) gyri; 4C) the association cortical thickness and SH retrieval accuracy for specific age group(s) for which there was a significant conditional mediation effect.

The bootstrap results indicated left angular gyrus cortical thickness significantly mediated the effect of age on SH retrieval accuracy only at ages 24 (Lower CI to Upper CI: 0.00 to 0.005) and 27 (Lower CI to Upper CI: 0.00 to 0.004), but not in later life. Post-hoc within age-group simple regression analyses of ROI→ performance confirmed a significant association between left angular gyrus cortical thickness and SH retrieval accuracy in young adults (F(1, 38) = 5.99, p =0.02). Figure 4 presents scatterplots showing the association between age and left angular gyrus cortical thickness (4A, ii), the association between left angular gyrus cortical thickness and SH retrieval accuracy across all ages (3B, ii), and the association between left angular gyrus cortical thickness and SH retrieval accuracy in young adults (aged 18 –35yrs) (4C, ii). From these graphs it appeared that one young adult (circled in red), may have been driving this conditional mediation effect. Indeed when we excluded this participant and re-ran the conditional mediation model in the left hemisphere, the conditional mediation effect in left lingual gyrus remained significant at ages 24 (Lower CI to Upper CI: −0.002 to −0.0004), 27 (Lower CI to Upper CI: −0.0037 to −0.0004) and 48 (Lower CI to Upper CI: −0.0027 to −0.0005), but the effect in left angular gyrus was no longer significant at ages 24 (Lower CI to Upper CI: −0.0002 to +0.005) and 27 (Lower CI to Upper CI: −0.0001 to +0.0047). Interestingly, after accounting for the conditional mediation effect of lingual gyrus cortical thickness on SH retrieval accuracy, the effect of age on SH retrieval accuracy was not significant in this model (Age, t = −1.66, p > 0.05).

## 4. Discussion

The current study used a cross-sectional design to examine the relationships between age, cortical thickness and episodic memory, as measured by spatial context memory tasks, across the adult lifespan. The behavioral data indicated that there was a significant age-related decline in both easy and hard versions of a spatial context memory task across the adult lifespan, which is consistent with prior lifespan studies of spatial context memory (Cansino et al., 2015). The cortical thickness analysis identified age-associated declines in bilateral superior frontal gyrus (SFG); caudal middle frontal gyrus (MFG); inferior frontal gyrus (IFG); supramarginal gyrus; angular gyrus; lingual gyrus; fusiform gyrus; gyrus rectus; and parahippocampal gyrus (PHG). These results are consistent with the meta-analysis of six studies (N = 883) conducted by Fjell et al. (2009), in which they reported a distributed pattern of age-related declines in cortical thickness in frontal, temporal, parietal and occipital gyri. The current study found that age-related differences on hard versions of the spatial context memory task (SH retrieval accuracy) was mediated or conditionally mediated by 3 brain regions: right superior frontal gyrus (SFG), right caudal middle frontal gyrus (MFG), and left lingual gyrus. In addition, right PHG cortical thickness significantly predicted SH retrieval accuracy, independent of age but did not significantly mediate the effect of age on SH retrieval accuracy. Although there were age-related differences in the easy version of the spatial context memory task (SE), this age effect was not significantly mediated or conditionally mediated by the cortical thickness of the ROIs examined in the current study.

Previous studies have examined how regional cortical volumes or thickness relates to episodic memory and generally identified several regions, not just five regions, which predicted memory across the adult lifespan (Kalpouzos, Chetelat, Baron, et al., 2009; Walhovd et al., 2006; Walhovd et al., 2004). This discrepancy in results is related to the analytic methods used. Past studies have generally employed multivariate general linear models and controlled for age, biological sex, and other measures (i.e. IQ or education) to examine how cortical thickness/volumes in the brain was associated to memory function. However, the current study used a more restrictive approach wherein mediation and conditional mediation was directly tested simultaneously across all ROIs, while taking into account age and biological sex. Thus, our results show that cortical thickness in right SFG, right MFG, and left lingual gyrus mediated, or conditionally mediated the effect of age on SH retrieval accuracy. Indeed, in our mediation models, after accounting for these ROI mediational affects, age was no longer significantly predictive of memory. In the following sections we discuss our regression result for right PHG and then the significant mediation and conditional mediation effects observed in left lingual gyrus, right MFG, and right SFG.

### 4.1. Right PHG cortical thickness and spatial context memory

We found that RPHG cortical thickness predicted SH retrieval accuracy, after controlling for age. This effect was marginally significant (p =0.052). Moreover, right PHG cortical thickness did not mediate nor conditionally mediate the effect of age on spatial context memory. The fact that the association we observed involved the right, not the left, PHG may be due the fact that we employed face stimuli and focused on spatial context memory. Indeed, lesion studies have demonstrated that patients with specific lesions to the right parahippocampal cortex are impaired on spatial memory tasks (Veronique D. Bohbot et al., 1998; Véronique D. Bohbot et al., 2000; Ploner et al., 2000; Weniger & Irle, 2006). Also, the lateralization of encoding activity in medial temporal cortex has been related to stimulus type: left activity has been associated with word encoding, bilateral with picture encoding, and right with face encoding (Golby et al., 2001; Powell et al., 2005). Therefore, individual differences in right PHG cortical thickness predicted spatial context memory, independent of age; but age-related declines in spatial context memory were not mediated by thinner right PHG cortex with advanced age. Instead, age-related differences in spatial context memory were mediated by other brain regions, i.e. the lingual and frontal gyri, discussed below.

### 4.2. Left lingual cortical thickness mediates age-related differences in spatial context memory at midlife

In the current study left lingual gyrus was the only brain region in which cortical thickness was found to mediate age-related differences in SH retrieval accuracy *across the adult lifespan*.

Specifically, there was a positive association between left lingual cortical thickness and SH retrieval accuracy across the adult lifespan.

Prior studies have also identified a positive association between gray matter volumes or cortical thickness in lingual gyrus and memory performance. Kalpouzos et al (2008) reported a significant positive association between gray matter volume in lingual gyrus and other brain regions, measured using voxel-based morphometry (VBM), and subsequent item memory for verbal stimuli in young, middle-aged and older adults. Walhovd et al (2006) also reported a positive association between cortical thickness in a variety of brain regions, including lingual gyrus, and higher verbal recall across months in an adult lifespan sample. More recently, functional MRI studies have highlighted the importance of ventral visual stream, which include the lingual gyrus, in supporting vivid encoding and contextual recollection (Morcom, 2014; Robinson-Long, Eslinger, Wang, Meadowcroft, & Yang, 2009; Skinner, Grady, & Fernandes, 2010). Indeed it has been suggested that successful episodic recollection relies on the recapitulation of the encoding context at retrieval, particularly in the occipital regions (Kahn, Davachi, & Wagner, 2004; McDonough, Cervantes, Gray, & Gallo, 2014; Morcom, 2014; St-Laurent, Abdi, Bondad, & Buchsbaum, 2014; Wang, Johnson, de Chastelaine, Donley, & Rugg, 2016). For example, using representational similarity analysis Wing et al. found that the similar occipital cortex activity during encoding and retrieval predicted memory success for specific items (Wing, Ritchey, & Cabeza, 2015). Age-related declines in the reactivation of occipital regions from encoding to retrieval have been associated with episodic memory deficits, particularly for recollection of contextual/associative information, with age (Dennis, Bowman, & Peterson, 2014; McDonough et al., 2014; St-Laurent et al., 2014).

Interestingly, in the current study a significant conditional mediation effect of left lingual cortical thickness on age-related differences in SH retrieval accuracy was only observed in young and middle-aged adults. This suggests that age-related declines in SH retrieval accuracy at these stages in adult development are specifically mediated by differences in left lingual gyrus thickness. However, the scatterplot and pot-hoc within group regression analyses suggest that thickness in this region was more strongly related to memory success in middle aged adults, and to a lesser degree in young adults. This is intriguing since prior fMRI adult lifespan studies of episodic memory have consistently reported altered ventral visual stream activity in middle-aged adults, compared to young adults, at encoding and retrieval (Grady, Springer, Hongwanishkul, McIntosh, & Winocur, 2006; H. Park, Kennedy, Rodrigue, Hebrank, & Park, 2013; J. Park et al., 2012). Moreover, we have found that context memory decline in midlife was associated with deficits in occipital activity at retrieval (E. Ankudowich, Pasvanis, S., Rajah, M. N., 2016; Kwon et al., 2016). Therefore, the current results build on prior findings by showing that left lingual cortical thickness may mediate memory success for both item *and associative memory*, and that midlife is a critical time in adulthood when maintenance of occipital cortex structure/function may be especially important for maintaining episodic memory for contextual details.

### 4.2. Frontal cortical thickness mediates age-related differences in context memory at midlife and older age

Right SFG cortical thickness conditionally mediated the effect of age on SH retrieval accuracy in middle-aged and older adults, but not young adults; and, right MFG cortical thickness conditionally mediated the effect of age on SH retrieval accuracy only in older adults. Earlier works have indicated that age-related declines in frontal volumes mediate deficits in cognitive control with age, which in turn impacts episodic memory performance (Head et al., 2008; Kalpouzos, Chetelat, Landeau, et al., 2009; Rajah, Languay, & Grady, 2011). Similarly, functional imaging studies have consistently reported right SFG and MFG activity during episodic memory tasks (Nyberg, Cabeza, & Tulving, 1996; Simons, Owen, Fletcher, & Burgess, 2005; Turriziani et al., 2008), particularly during the retrieval of contextual details (Cansino, Maquet, Dolan, & Rugg, 2002; Rajah & McIntosh, 2006; Ranganath, Heller, & Wilding, 2007; Rugg, Fletcher, Chua, & Dolan, 1999); and found that age-related declines in context memory were associated with functional and structural declines in right frontal cortex, particularly in right MFG (Cabeza, Anderson, Houle, Mangels, & Nyberg, 2000; de Chastelaine, Mattson, Wang, Donley, & Rugg, 2016; Dulas & Duarte, 2014; Maillet & Rajah, 2013; Rajah & D’Esposito, 2005; Rajah et al., 2011; Rajah, Languay, et al., 2010).

Interestingly in the current study the conditional mediation model indicated that right SFG and MFG exhibited opposite mediational influences on SH retrieval accuracy only in older age, such that having thicker right SFG and thinner right MFG predicted better SH retrieval accuracy in older adults. In a previous fMRI study of context memory in young and older adults, we observed that older adults exhibited functional declines in right MFG, and greater activity in right SFG. Moreover, in this study greater right SFG activity in older adults was correlated with better context retrieval. The current results provide parallel results in that thicker right SFG was associated with better SH retrieval accuracy in middle aged and older adults, but thicker right MFG was associated with worse SH retrieval accuracy in older adults. One speculative interpretation of the current findings is that relying on right MFG related processes, compared to SFG related processes, in later life can be detrimental to SH retrieval accuracy. Therefore, our result that age-related declines in spatial context memory are mediated by cortical thickness in right SFG and MFG in later life is consistent with the frontal lobe hypothesis of age-related declines in episodic memory (Spencer & Raz, 1994; Stuss, Craik, Sayer, Franchi, & Alexander, 1996; West, 1996). In addition, our results in older adults support the hypothesis that successful context memory in late life may be particularly dependent on the integrity of the right SFG (Dulas & Duarte, 2012; Kragel & Polyn, 2015; Rajah, Languay, et al., 2010).

However, there is also evidence that in older age, successful episodic memory is positively correlated to frontal activity (Cabeza, Anderson, Locantore, & McIntosh, 2002; Dennis & Cabeza, 2012; Rajah, Languay, et al., 2010). The PASA model suggests that older adults may increase activity in anterior frontal cortices to compensate for deficits in posterior cortical regions, i.e. the medial temporal lobes and sensory cortices, during memory tasks (Davis et al., 2008). Our current results are not entirely consistent with the PASA since right PHG and left lingual cortical thickness was positively associated with SH retrieval accuracy *across the adult lifespan*. However, left lingual cortical thickness mediated SH retrieval accuracy to a greater extent in young and midlife, and cortical thickness in right SFG positively mediated SH retrieval accuracy only in middle-aged and older adults. These results are consistent with the PASA model. Yet, we observed a negative mediation of SH retrieval accuracy by right MFG cortical thickness in older age. It is unclear if this result is consistent with PASA since it remains unknown how this difference in cortical thickness may relate to activity. Overall, the fact that frontal cortical thickness mediated SH retrieval accuracy with advanced age is generally consistent with the PASA model.

### 4.4. Limitations

The cross-sectional, as opposed to longitudinal design of the current study limited the interpretation of the results. A longitudinal study would have enabled us to examine intra-individual changes in cortical thickness and memory performance with age. This would have allowed us to make direct interpretations about how the brain ages, and how this affects changes in context memory performance. As the current study stands, inter-individuals differences are only suggestive of these changes.

Also, the current analysis only uncovered significant models when examining the difficult version of the spatial context memory task. We attributed this finding to the greater age-associated variability of the more difficult spatial hard task. It is possible that given a larger sample size, the models examining the easier spatial tasks would uncover similar results. Alternatively, it is possible that the current findings are not specific to spatial context memory, but more generalizable to being the result of increased difficulty. This remains unclear, as the current study only implemented spatial context memory tasks. Rather, it would be useful to analyze additional cognitive tasks at varying levels of difficulty in future study.

## 5. Conclusions

The current study sought to examine whether and how regional cortical thickness mediates age-related differences in spatial context memory. We found that age-related differences in context memory performance were significantly accounted for by cortical thickness of the right PHG independent of age, left lingual gyri among young and middle-aged adults, and, right caudal MFG and SFG among middle-aged and older adults. This pattern of results reflects a posterior to anterior shift with age in the structural neural correlates accounting for age-associated differences in memory performance. However, our results also suggest that some of the anterior, frontal, associations with SH retrieval accuracy were negative in late life; and that there are dissociation in right frontal contributions to successful contextual retrieval in later life.

## Acknowledgements

This work was supported by the Canadian Institutes of Health Research (CIHR) operating grant (Grant No. 126105) and the Alzheimer’s Society of Canada (Grant No. 1435) awarded to MNR. We thank R. Patel for help with conducting the CIVET analysis.

---

**Table S1.** Path coefficients for spatial hard accuracy conditional mediation model with right hemisphere ROIs.

| **Paths** | **Coefficients** |
| --- | --- |
| Age → right parahippocampal gyrus | -0.0045*** |
| Age → right gyrus rectus | -0.0042*** |
| Age → right superior frontal gyrus | -0.0047*** |
| Age → right caudal middle frontal gyrus | -0.0053*** |
| Age → right inferior frontal gyrus | -0.0053*** |
| Age → right middle temporal gyrus | -0.0040*** |
| Age → right angular gyrus | -0.0037*** |
| Age → right supramarginal gyrus | -0.0051*** |
| Age → right superior temporal gyrus | -0.0066*** |
| Right parahippocampal gyrus → spatial hard accuracy | 0.50* |
| Right gyrus rectus → spatial hard accuracy | 0.26 |
| Right angular gyrus → spatial hard accuracy | -0.42 |
| Right supramarginal gyrus → spatial hard accuracy | -0.25 |
| Right superior temporal gyrus → spatial hard accuracy | -0.473 |
| Right middle temporal gyrus → spatial hard accuracy | 0.58 |
| Right superior frontal gyrus → spatial hard accuracy | -0.84* |
| Right caudal middle frontal gyrus → spatial hard accuracy | 0.45 |
| Right inferior frontal gyrus → spatial hard accuracy | 0.11 |
| Age*right parahippocampal gyrus → spatial hard accuracy | -0.009 |
| Age*right gyrus rectus → spatial hard accuracy | -0.006 |
| Age*right angular gyrus → spatial hard accuracy | 0.007 |
| Age*right supramarginal gyrus → spatial hard accuracy | 0.009 |
| Age*right superior temporal gyrus → spatial hard accuracy | 0.008 |
| Age*right middle temporal gyrus → spatial hard accuracy | -0.014 |
| Age*right superior frontal gyrus → spatial hard accuracy | 0.026** |
| Age*right middle caudal frontal gyrus → spatial hard accuracy | -0.015 |
| Age*right inferior frontal gyrus → spatial hard accuracy | 0.0004 |
| Age → spatial hard accuracy | -0.017 |

\*p<0.05; \*\*p<0.01; \*\*\*p<0.001
a. The spatial easy accuracy conditional mediation model was non-significant at $p > 0.05$ .

**Table S2.** Path coefficients for spatial hard accuracy conditional mediation model with left hemisphere ROIs.

| **Paths** | **Coefficients** |
| --- | --- |
| Age → left parahippocampal gyrus | -0.0048*** |
| Age → left fusiform gyrus | -0.0028*** |
| Age → left gyrus rectus | -0.0037*** |
| Age → left superior frontal gyrus | -0.0053*** |
| Age → left caudal middle frontal gyrus | -0.0065*** |
| Age → left inferior frontal gyrus | -0.0062*** |
| Age → left supramarginal gyrus | -0.0052*** |
| Age → left angular gyrus | -0.0035*** |
| Age → left lingual gyrus | -0.0045*** |
| Left angular gyrus → spatial hard accuracy | -1.02 |
| Left lingual gyrus → spatial hard accuracy | 0.56 |
| Left fusiform gyrus → spatial hard accuracy | -0.56 |
| Left gyrus rectus → spatial hard accuracy | 0.62 |
| Left parahippocampal gyrus → spatial hard accuracy | 0.16 |
| Left superior frontal gyrus → spatial hard accuracy | -0.26 |
| Left caudal middle frontal gyrus → spatial hard accuracy | 0.49 |
| Left inferior frontal gyrus → spatial hard accuracy | -0.13 |
| Left supramarginal gyrus → spatial hard accuracy | -0.06 |
| Age*left lingual gyrus → spatial hard accuracy | -0.005 |
| Age*left fusiform gyrus → spatial hard accuracy | 0.007 |
| Age*left gyrus rectus → spatial hard accuracy | -0.011 |
| Age*left parahippocampal gyrus → spatial hard accuracy | -0.0034 |
| Age*left superior frontal gyrus → spatial hard accuracy | 0.007 |
| Age*left caudal middle frontal gyrus → spatial hard accuracy | -0.011 |
| Age*left inferior frontal gyrus → spatial hard accuracy | 0.0038 |
| Age*left angular gyrus → spatial hard accuracy | 0.017 |
| Age*left supramarginal gyrus → spatial hard accuracy | -0.033 |
| Age → spatial hard accuracy | -0.031 |

\*p<0.05; \*\*p<0.01; \*\*\*p<0.001
a. The spatial easy accuracy conditional mediation model was non-significant at $p > 0.05$ .

